# Novel metabolic role for CD47 in pancreatic β-cell insulin secretion and islet transplant outcomes

**DOI:** 10.1101/2022.07.22.501201

**Authors:** Kedar Ghimire, Atharva Kale, Jennifer Li, Sohel M. Julovi, Philip O’Connell, Shane T. Grey, Wayne J. Hawthorne, Jenny E. Gunton, Natasha M. Rogers

## Abstract

Diabetes is a global public health burden and is characterized clinically by a relative or absolute insulin deficiency. Therapeutic agents that stimulate and improve insulin secretion and insulin sensitivity are in high demand as diabetic treatment. CD47 is a cell surface glycoprotein implicated in multiple cellular functions, including recognition of self, angiogenesis, and nitric oxide signaling, however its role in the regulation of insulin secretion remains unknown. For the first time we demonstrate that CD47 receptor signaling inhibits insulin release from β-cells and that it can be pharmacologically exploited to boost insulin secretion. CD47 depletion stimulates insulin granule exocytosis via activation of the Rho GTPase Cdc42. CD47 deficiency improved glucose clearance and insulin sensitivity in mice. CD47 blockade enhanced islet transplantation efficiency and improved outcomes. Further, anti-CD47 antibody treatment delayed the onset of diabetes in non-obese diabetic mice and protected them from overt diabetes. Our findings identify CD47 as a previously unrecognized regulator of insulin secretion and its manipulation in β-cells offers a novel therapeutic opportunity for diabetes and islet transplantation by correcting insulin deficiency.

**One Sentence Summary:** CD47 limits insulin secretion and islet transplant outcomes

## INTRODUCTION

Insulin is crucial for cellular glucose uptake, as well as regulating carbohydrate, lipid, and protein metabolism. Failure to produce sufficient insulin due to pancreatic ß-cell dysfunction and a loss of β-cell mass are features of both type 1 (T1DM) and type 2 diabetes mellitus (T2DM). T1DM leads to near-absolute insulin deficiency through β-cell destruction by autoreactive T cells [1], affecting β-cell population that is poorly equipped to survive an inflammatory microenvironment. Patients with obesity respond to insulin resistance with insulin hypersecretion and expansion of β-cell mass, leading to hyperglycemia when β-cells are no longer able to compensate [2]. Islet transplantation is a successful, routine clinical procedure currently available for select patients with unstable T1DM and hypoglycemia unawareness [3]. However, a limited supply of donor pancreases suitable for islet isolation, as well as significant β-cell attrition and limited insulin secretory capacity in the peri-transplant period has meant restricted applicability for this treatment. Efforts to stimulate insulin secretory capacity on one side while improving insulin sensitization may lead to additional therapeutic opportunities for diabetes and improve islet transplant outcomes.

Insulin secretion is exquisitely tuned to metabolic demand, particularly plasma glucose concentration, through a process that is facilitated by ß-cell proximity to the pancreatic vascular network [4]. Under physiological conditions, blood glucose is maintained within a narrow range, coordinated by nutrient availability and a biphasic glucose-responsive insulin secretion from β-cells [5]. The transient first phase triggers immediate exocytosis of insulin from pre-docked granules at the plasma membrane, and the second phase requires granule mobilization from an intracellular pool to the membrane for sustained insulin release [6]. Actin remodeling facilitates the mobilization of insulin granules from the Golgi complex to the cell membrane, and the Rho GTPase protein Cdc42 is a key component of cytoskeletal rearrangement [7]. Cdc42 has been implicated as a proximal mediator of insulin release, and dysfunction of Cdc42 impairs insulin secretion [8]. The Src family of protein tyrosine kinases, specifically Yes kinase, activates Cdc42 and provides a proximal step in glucose-specific release of insulin [9]. Similarly, pharmacological Lck/Yes novel (Lyn) kinase activation has been implicated in insulin receptor potentiation and elicited both rapid-onset and durable improvement in glucose homeostasis in animal models of diabetes [10]. Upstream regulators of these kinases or Cdc42 in β-cells remain elusive.

CD47 is an integral membrane glycoprotein with universal cellular expression. By engaging the high affinity ligand thrombospondin-1 [11], cross-talk with signal inhibitory regulatory protein (SIRP) [12], or specific association with integrins [13], CD47 participates in a broad range of cellular functions, including recognition of self [14], nitric oxide signaling [15], autophagy [16], and oxidative stress [17]. Recent kinetic studies have demonstrated that CD47 can cluster in lipid rafts on the cell surface which are enriched for kinases capable of executing signal transduction [18]. The role of CD47 signaling in β-cell function remains unexplored. Here we demonstrate for the first time that CD47 suppresses insulin secretion by deactivating Cdc42, and inhibition of CD47 in islets enhances β-cell function. In addition, CD47 expression limits islet transplant outcomes and treatment of non-obese diabetic mice with a CD47 blocking antibody delays the onset of overt diabetes. Our findings have direct relevance for the treatment of T1DM by prolongating insulin independence, T2DM as an adjuvant therapeutic, and islet transplantation to enhance efficacy of islets derived from deceased donors.

## RESULTS

### CD47 is preferentially expressed by pancreatic islets

Our previous work has demonstrated superior glucose clearance and enhanced insulin sensitivity in aged (18+ months) CD47-/- mice compared to littermate controls [19]. We examined the relevance of these findings to pancreatic endocrine functions. With immunofluorescence, we observed that CD47 was preferentially expressed in murine pancreatic islets compared to surrounding acinar tissue (Fig. 1a-b). Human islets also expressed CD47 which co-localized with insulin expression in β-cells (Fig. 1c). CD47 was expressed by MIN-6 cells (Fig. 1d), a widely used β-cell line of murine origin that has been experimentally validated to represent key physiological processes is islets, including insulin secretion, glycolysis, and oxidative phosphorylation [20]. The conserved expression of endogenous CD47 in human, mouse and cultured β-cell models suggests a potential physiological role for CD47 in these cells.

**Figure 1.**
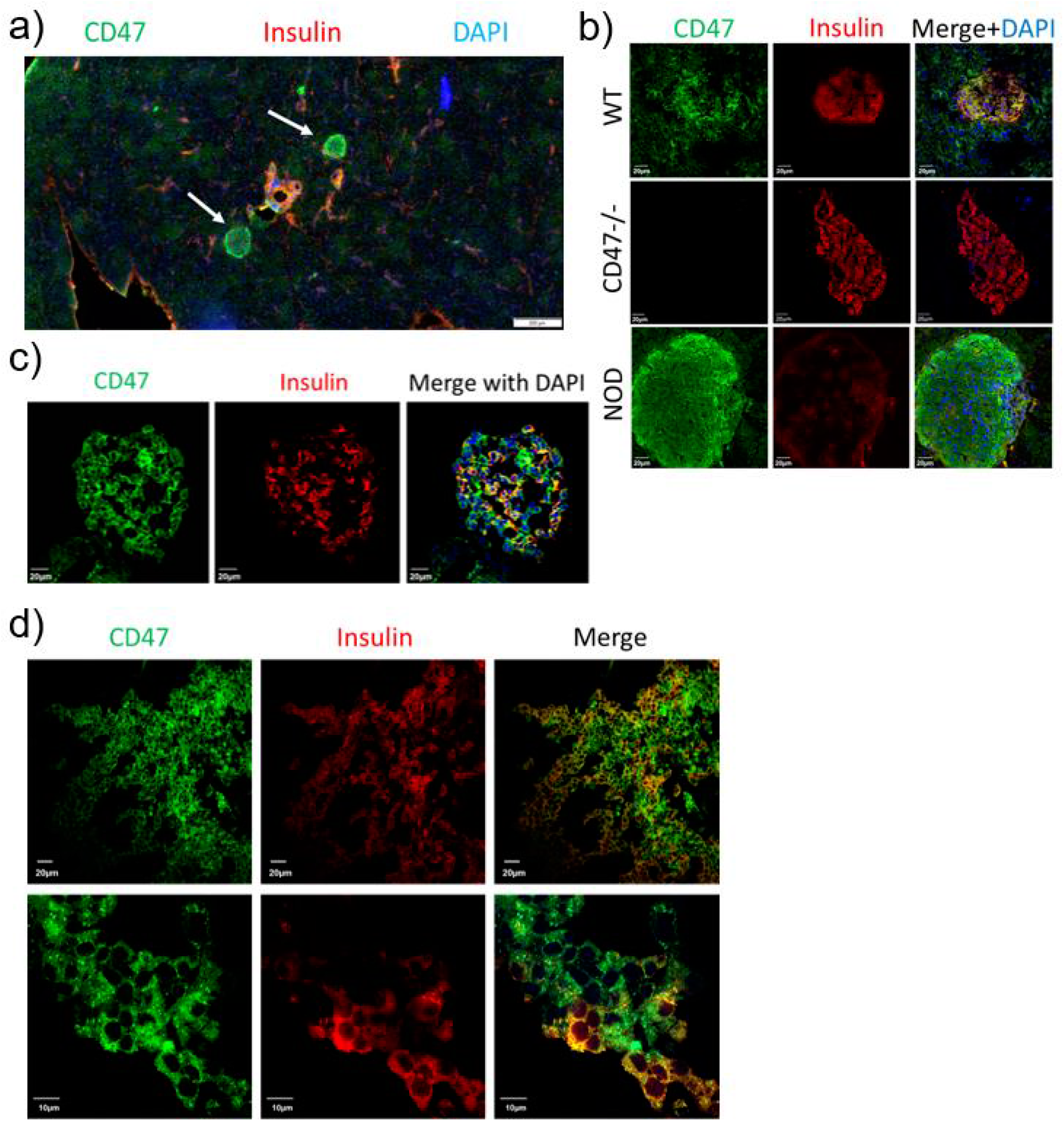
CD47 is preferentially expressed in pancreatic islets. Immunofluorescent staining of CD47 (green) and insulin (red) in a) mouse pancreas (islets of Langerhans indicated by white arrows), scale bar: 200 um, b) wild-type (WT), CD47^-/-^ and NOD islets, c) human islets, scale bar all panels: 20 um, and d) MIN6 cells, scale bars: upper panel (20 um), lower panel (10 um).

### CD47 impairs glucose-stimulated insulin secretion from mice and human pancreatic β-cells

Pancreatic β-cells reside in a complex microenvironment, where they interact with other endocrine cells, as well as vascular endothelium and immune cells [21]. We, and others, have previously reported that CD47 promotes deterioration of several homeostatic mechanisms, including angiogenesis and blood flow [22, 23]. CD47 expression increased in the vasculature of aged mice and humans, and aged CD47^-/-^ mice were protected from high fat diet (HFD)-induced weight gain [19]. It is not known whether CD47 expression changes during insulin resistant states and whether this is involved in regulating insulin release and glycemic control. To investigate whether CD47 receptors in β-cells are regulated depending on energy status, we isolated pancreatic β-cells (dithizone-positive) from lean and 16-week HFD-fed WT (CD47^+/+^) mice. Notably, CD47 expression was significantly downregulated in obese hyperinsulinemic mice (Fig. 2a) suggesting that reduced CD47 receptor signaling may be a compensatory mechanism to allow β-cells to secrete more insulin in the context of insulin resistance.

**Figure 2.**
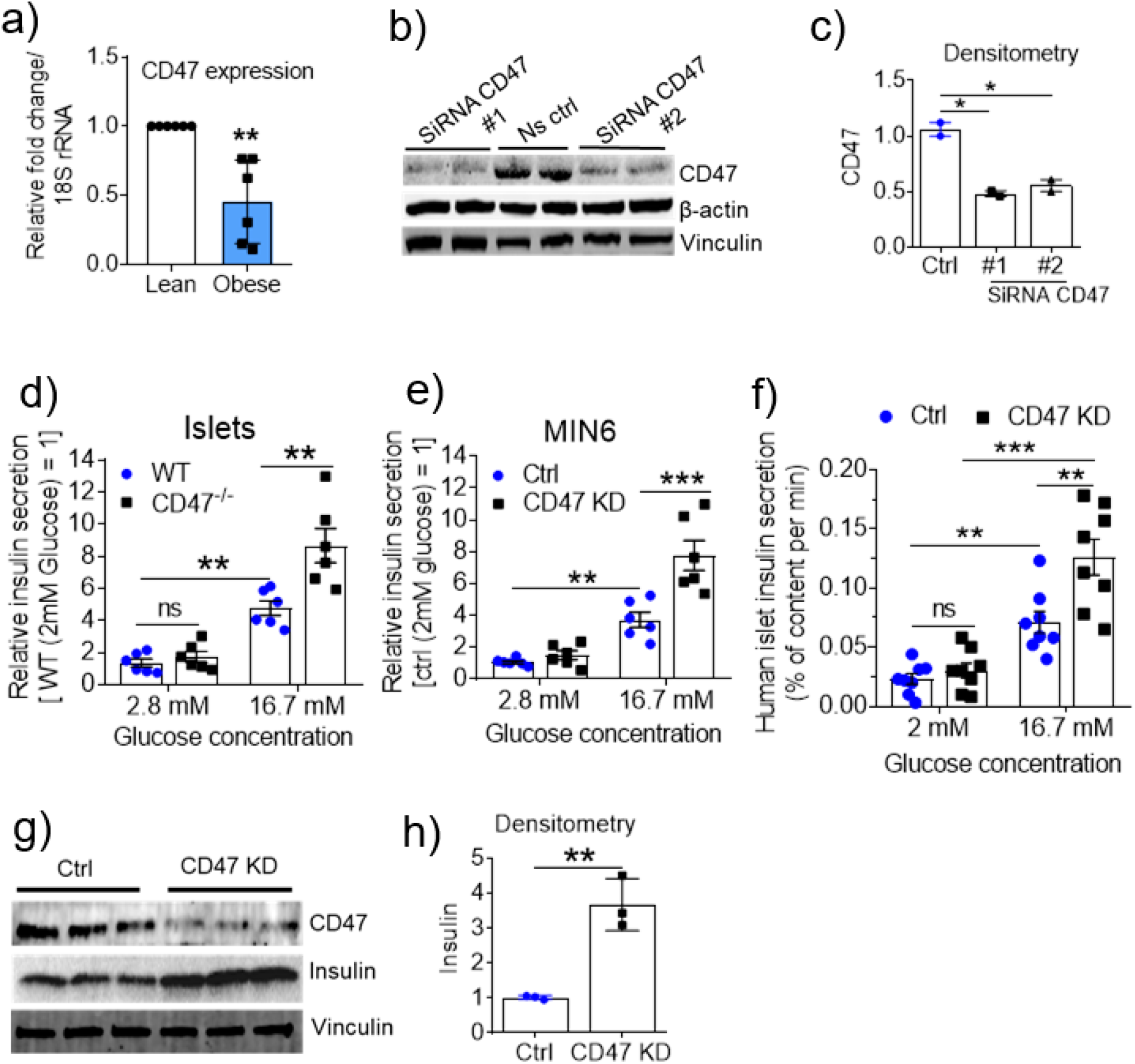
CD47 impairs glucose-stimulated insulin secretion from mice and human pancreatic β-cells. a) CD47 receptor mRNA expression in primary murine β-cells isolated from 16 week old lean or obese WT mice, determined by qRT-PCR. b) Primary murine β-cells were transfected with non-silencing control (Ctrl) or CD47 siRNA, then probed for CD47 and representative western blot analysis and c) quantification of knock-down efficiency. d) Glucose-stimulated insulin secretion (GSIS) in β-cells isolated from wild-type (WT, CD47+/+) or CD47^-/-^ mice. e) GSIS in MIN6 cell line following transfection with control (Ctrl) or knockdown (CD47 KD). f) GSIS in isolated human islets (n=8 donors). g) Representative western blot analysis and (h) quantification of insulin expression following control (ctrl) and CD47-knocked down (CD47 KD) cells. Conditions are as indicated in figure panels with quantification of changes in insulin secretion levels relative to islets/cells treated with 2.8 mM glucose. Data are presented as mean ± SEM from triplicate wells from n=3 independent experiments; *p<0.05, **p<0.01, ***p<0.001 by t-test (b, g) or 2-way ANOVA (c,d,e) followed by Tukey’s multiple comparisons test.

CD47 could be readily detected and manipulated with RNAi (>50% knockdown) in isolated primary mouse islets (Fig 2b, c). We then performed glucose-stimulated insulin secretion (GSIS) experiments on islets isolated from WT or CD47^-/-^ mice. As shown in Fig. 2d, CD47^-/-^ islets showed no significant difference in insulin secretion under basal glucose conditions (2.8 mM), but insulin secretion was significantly increased following exposure to high (16.7 mM) glucose conditions. Similar responses were observed in the MIN6 cells (Fig. 2e) and isolated human islets following CD47 knockdown (Fig. 2f). We further examined if CD47 signaling limited insulin content in β-cells under high glucose conditions. Silencing of CD47 led to upregulation of insulin expression within MIN6 cells, detected by western blotting (Fig. 2g, h). Together, these data demonstrate a previously unknown role for CD47 receptor signaling as a negative regulator of insulin release in pancreatic β-cells under a pathophysiological glucose load.

### CD47 limits the *in vivo* response to a glucose load

In previous studies from our group, we observed improved glucose tolerance in aged CD47^-/-^mice compared to WT controls [19]. We next examined whether CD47 plays a role in β-cell physiology in young mice. To test this, CD47^-/-^ mice were subjected to an intraperitoneal glucose tolerance test (IPGTT). CD47^-/-^ mice showed superior glucose tolerance overall (Suppl. Fig. 1a), associated with improved insulin secretion. Serum insulin levels were significantly different between the two genotypes following a glucose load (Fig. 3a, b). In a separate cohort, we compared insulin sensitivity between WT and CD47^-/-^ mice. Insulin tolerance tests revealed an enhanced insulin response in CD47^-/-^ mice compared to controls (Fig 3c, d). These results demonstrate that glucose clearance and insulin sensitivity are significantly improved in CD47^-/-^ mice. Previous studies have demonstrated that Lyn kinase activation plays a crucial role in improving insulin sensitivity through insulin receptor potentiation, eliciting a rapid-onset and durable improvement in glucose homeostasis in animal models of type 2 diabetes [10]. Phosphorylation of Lyn at Tyrosine 396 (Y396) alters its structure and increases the kinase activity [24]. We examined whether CD47 loss-of-function affected Lyn phosphorylation in β-cells. Silencing of CD47 with siRNA, or antibody blockade of CD47, in MIN6 cells led to increased phosphorylation of Lyn kinase at the Y396 residue (Fig. 3e, f). These results show that insulin sensitivity was enhanced in CD47^-/-^ mice with an extended duration and magnitude of insulin response which could be mediated through activation of Lyn kinase.

**Figure 3.**
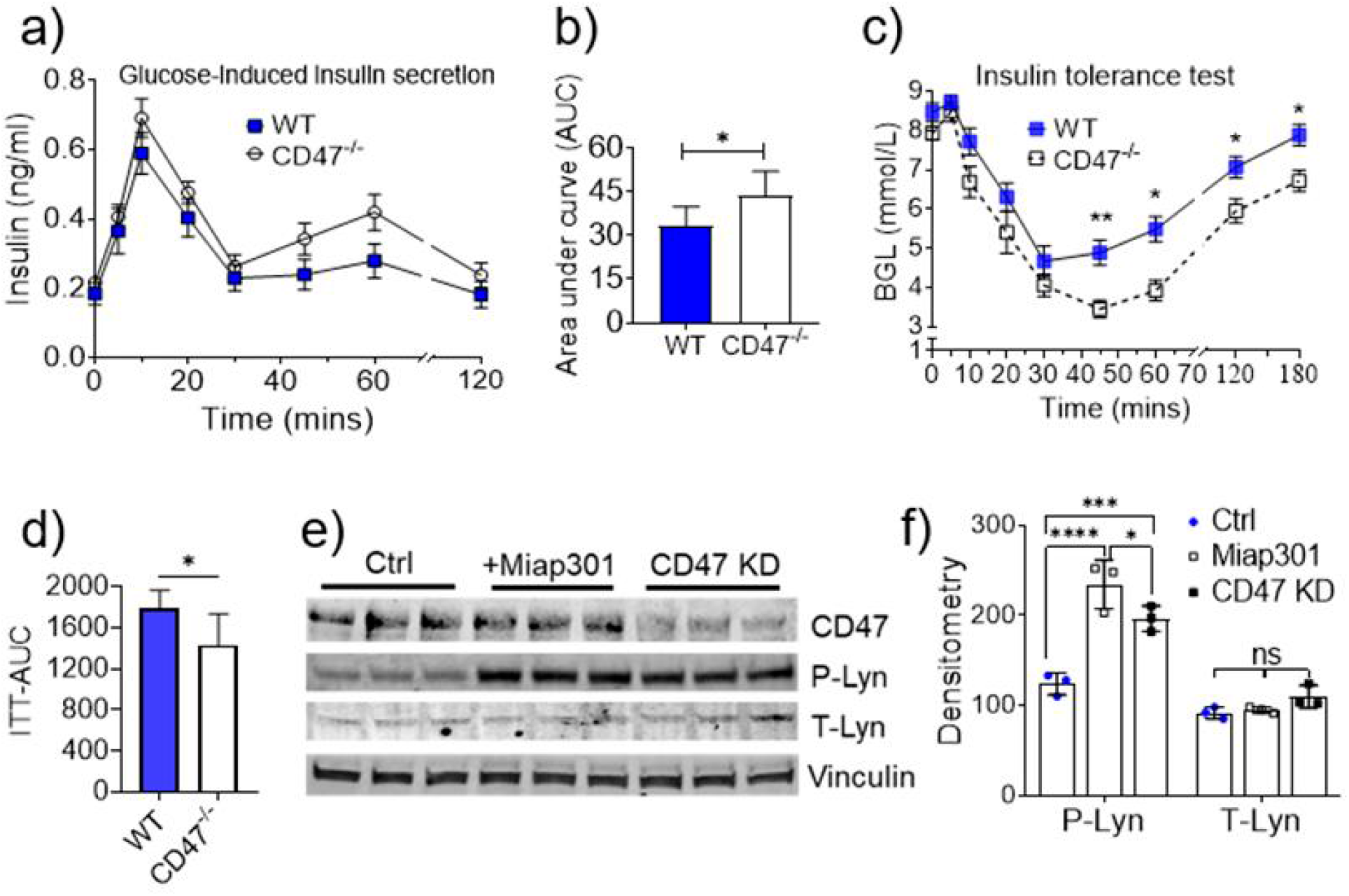
Mice lacking CD47 receptor have altered glucose metabolism. a) Serum insulin levels in CD47^-/-^ and WT mice after an oral glucose load, and b) quantification of area under the curve (AUC). c) Time course and (d) AUC of blood glucose concentration following an insulin tolerance test in WT and CD47^-/-^ mice. (e, f) MIN6 cells were treated with Miap301, or transfected with control (Ctrl) or CD47 siRNA (CD47 KO) and lysates probed for total and phospho-Lyn kinase. Representative western blot and combined densitometry are shown. Data are presented as mean ± SEM from n=7-8 mice per group or triplicate wells for cell lines, *p < 0.05, ** p < 0.01, **** p < 0.0001 by t-test (b, d) or 2-way ANOVA (f) followed by Tukey’s multiple comparisons test.

### CD47 signaling regulates insulin secretion via Cdc42 activation

Insulin granules are the hormone repository of the β-cell. To investigate if deletion of CD47 affected insulin granule secretion at nanomolecular resolution, glucose-stimulated insulin granule docking was examined using transmission electron microscopy (TEM). β-cells were isolated from WT or CD47^-/-^ pancreas, subjected to GSIS (exposure to 2.8- and 16.7-mM glucose concentrations) and samples processed for visualisation (Fig. 4a). We quantified the total number of insulin granules per surface area (μm^2^). At basal glucose levels, CD47^-/-^ β-cells did not demonstrate significantly higher insulin granule numbers compared to WT (Fig. 4b). After glucose stimulation (16.7 mM), both WT and CD47^-/-^ mice showed a significant reduction in surface area β-cell granules (Fig. 4b) but this decrease was markedly higher in WT than CD47^-/-^ β-cells (Fig. 4c). However, after glucose stimulation, a significantly greater number of granules were docked at the plasma membrane of CD47^-/-^ β-cells compared to WT (Fig. 4d). This was accompanied by a significant increase in the number of exocytosing granules in CD47^-/-^ β cells compared to WT (Fig. 4e, Suppl. Fig 2a, b). However, there was no difference in the insulin-containing dense core granule size between WT and CD47^-/-^ β-cells under either glucose concentration (Fig. 4f). These results show that CD47 deficiency facilitates both insulin granule docking and exocytosis.

**Figure 4.**
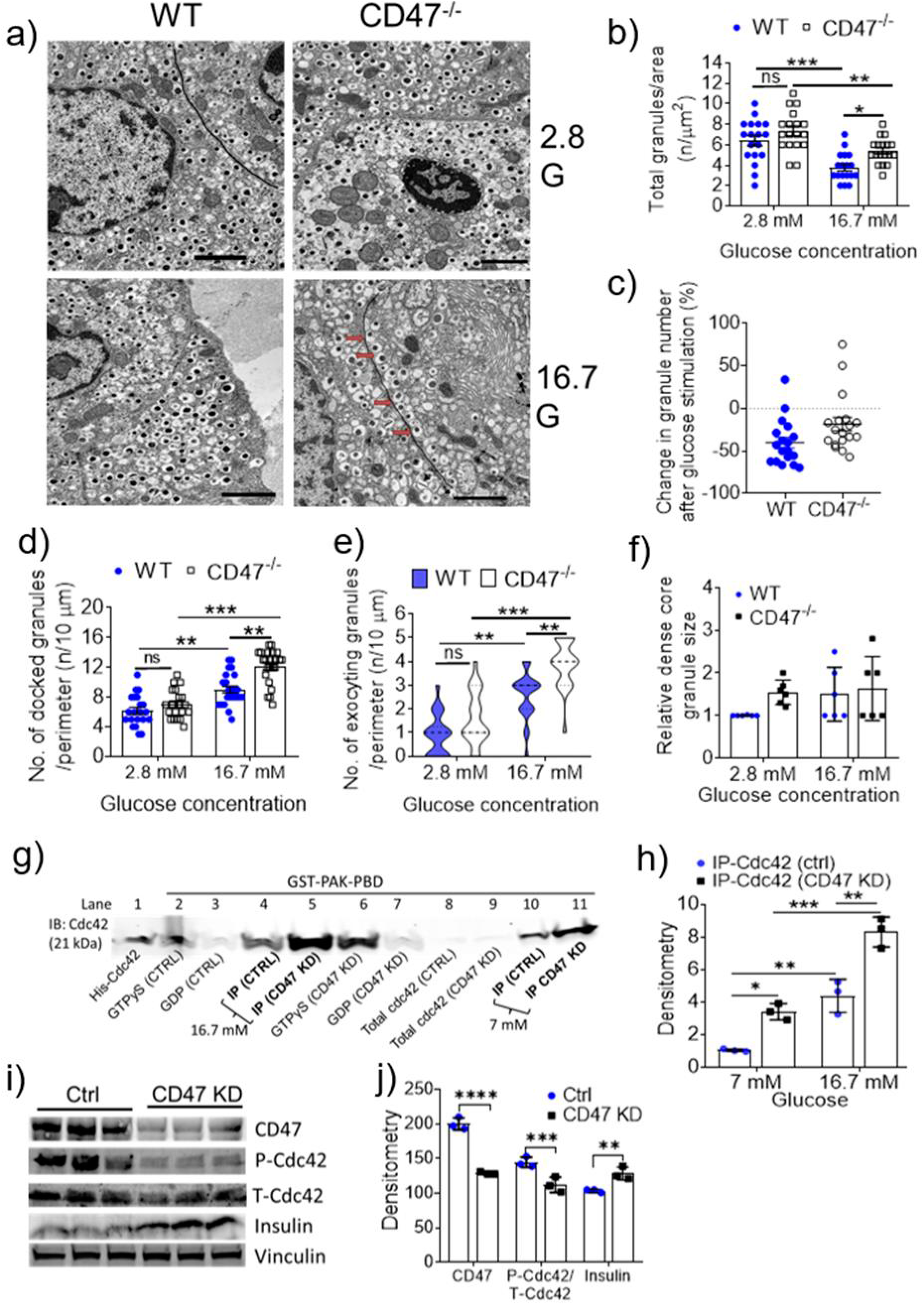
CD47 signaling limits insulin secretion in β-cells. a) Representative transmission electron micrograph images of β-cells from WT and CD47^-/-^ mice with glucose (G) concentrations as indicated. Scale bar=2 um. β-cells from WT and CD47-/- mice were assessed for (b) total insulin-containing vesicles, (c) percent change in granule number after glucose load, (d) quantification of docked vesicles (closer than 0.13 μm to the plasma membrane), e) quantification of vesicles that were at or releasing from the plasma membrane, and f) comparison of dense core granule size. β-cells transfected with control (Ctrl) or CD47 siRNA (CD47 KD) under low or high glucose conditions were probed for activated Cdc42 (GTP-bound) levels, (g) Representative western blot and h) densitometric quantification, i) Representative western blot and (j) quantification of Cdc42 phosphorylation levels under 16.7 mM glucose. Data are presented as mean ± SEM, *p < 0.05, ** p < 0.01, ***p < 0.001 by 2-way ANOVA followed by Tukey’s multiple comparisons test.

We then speculated whether CD47’s direct impact on docking and discharge of insulin granules was related to changes in cytoskeletal proteins that regulate the intracellular granule pool. Cdc42 signaling is essential for insulin secretion through actin remodeling and granule exocytosis, and dysfunctional Cdc42 impairs insulin secretion and contributes to diabetes [25–27]. However, upstream regulatory mechanisms of Cdc42-mediated insulin secretion have remained elusive.

We performed immuno-precipitation experiments to determine whether CD47 inhibition altered activation of Cdc42 in β-cells. Following a high glucose load (16.7 mM), activation of Cdc42 was higher in CD47-depleted β-cells compared to the controls (Fig. 4g, h; lanes 4 and 5). We also examined whether activation was dependent on glucose load by stimulating the cells with a lower glucose concentration (7 mM). This concentration of glucose stimulated insulin secretion, but not to the same degree, and the amount of activated Cdc42 (GTP-bound) precipitated was lower (Fig. 4g, lanes 10 and 11). Next, we investigated whether CD47 modulated phosphorylation of Cdc42. Phosphorylation is not a key event in activation/inactivation of Rho GTPases but it abrogates binding of Cdc42 to its downstream effectors such a N-WASP and Cofilin, thereby negatively affecting its activity [28]. Knockdown of CD47 inhibited Cdc42 phosphorylation at Ser-71 compared to its non-silencing control, simultaneously increasing insulin production in β-cells (Fig. 4i, j). These data demonstrate that CD47 impairs glucose-stimulated Cdc42 activation, thereby limiting insulin secretion.

### Enhanced islet transplantation efficiency utilizing CD47-deficient islets

We conducted a series of studies exploring the potential beneficial clinical effects of manipulating the CD47 receptor system in situations of limited β-cell mass. We performed islet transplantation experiments to investigate whether CD47 receptor-deficient islets could provide better metabolic control. We used streptozotocin (STZ) to induce DM in WT (C57BL/6) mice, and 1 week after STZ administration, 85% of the mice presented an average blood glucose of 28 mmol/L. No mice recovered spontaneously from STZ-induced diabetes. Islets of comparable size were hand-picked, and β-cell mass was calculated to ensure that material with equal insulin content was transplanted. As shown in Fg. 5a, β-cell mass was not significantly different between WT and CD47^-/-^ mice. Diabetic C57BL/6 mice were transplanted with equivalent numbers (400) of syngeneic islets from CD47^+/+^ (WT) or CD47^-/-^ mice, or a sufficient (optimal) mass of 550 WT islets to establish a baseline marker of islet quality. Blood glucose levels (BGL) were monitored for 20 days. As expected, mice receiving optimal islet mass showed immediate return of euglycemia (Fig. 5b), whereas mice receiving a minimal number of WT islets demonstrated only minor improvements in BGL by day 20 post-transplantation, which never completely normalized (Fig. 5b, c). In contrast, mice transplanted with 400 CD47^-/-^ islets showed significant glycemic improvement by day 8 post-transplantation, and by day 18, BGL were comparable with the mice receiving the optimal islet mass transplant (Fig. 5b, c). Consistent with the improved glycemic control, IPGTT performed at day 8 post-transplantation was also significantly improved in mice transplanted with CD47^-/-^ islets (Fig. 5d, e), in keeping with their improved insulin secretory response (Fig. 5f, g). Together, these data demonstrate that reduced CD47 receptor signaling in islets is advantageous for insulin secretory function, with direct relevance to clinical islet transplantation where donor islets are of limited supply.

**Figure 5.**
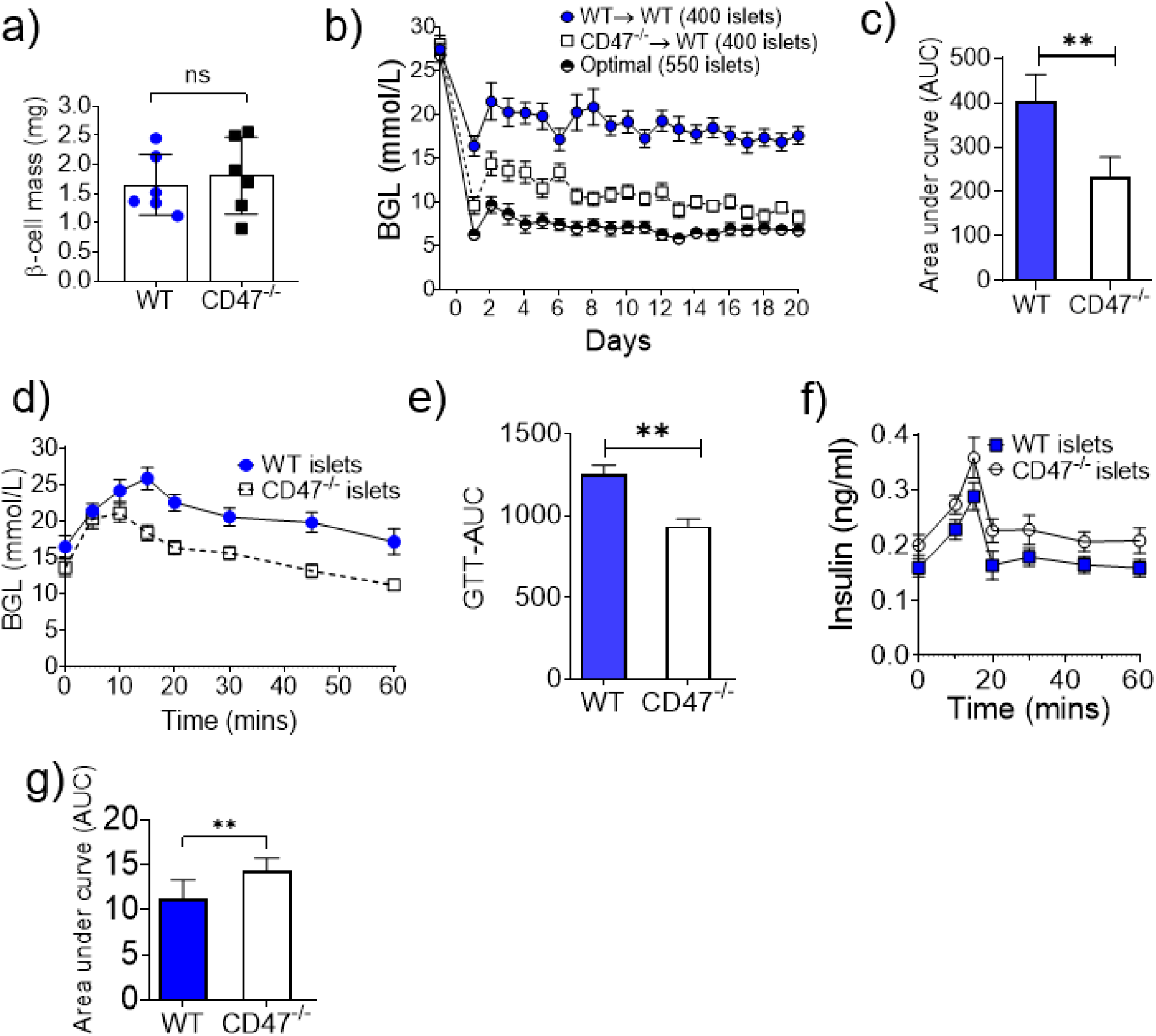
Improved glycemic control in diabetic mice transplanted with CD47^-/-^β-cells. a) Comparison of β-cell mass in 12 week-old WT and CD47^-/-^ mice (n = 6). Streptozotocin-induced diabetic mice were transplanted with an optimal number (550) of WT islets (n = 8), or a minimal number (400) of WT islets (n = 8) or CD47^-/-^ islets (n = 11). b) Time course and (c) quantification (area-under-the-curve, AUC) of post-transplant blood glucose levels (BGL). Transplanted mice (n=5) were fasted for 6 hours and intraperitoneal glucose injection (1 g/kg body weight) was performed on day 8 post-transplant. (d) BGL with (e) quantification (AUC) and (f) corresponding insulin levels with (g) quantification (AUC) were measured. Data are presented as mean ± SEM, *p < 0.05, ** p < 0.01 by unpaired t-test.

### Blockade of CD47 receptor signaling improves islet transplant outcomes

To test whether the beneficial effects of lack of CD47 signaling could be replicated pharmacologically, we examined the impact of Miap301- a functional blocking CD47 antibody on glucose-induced insulin secretion in vivo. A cohort of C57BL/6 mice were fasted and treated with Miap301 at a dose known to have physiological actions [29], 2 h prior to a bolus glucose load. Glucose clearance and serum insulin levels were then assessed. Mice that received Miap301 exhibited noticeable improvements in glucose clearance (Fig. 6a, b), with concurrently increased serum insulin levels (Fig. 6c, d), confirming that short-term blockade of CD47 receptor signaling can enhance physiologically-triggered insulin secretion and improve glucose homeostasis.

**Figure 6.**
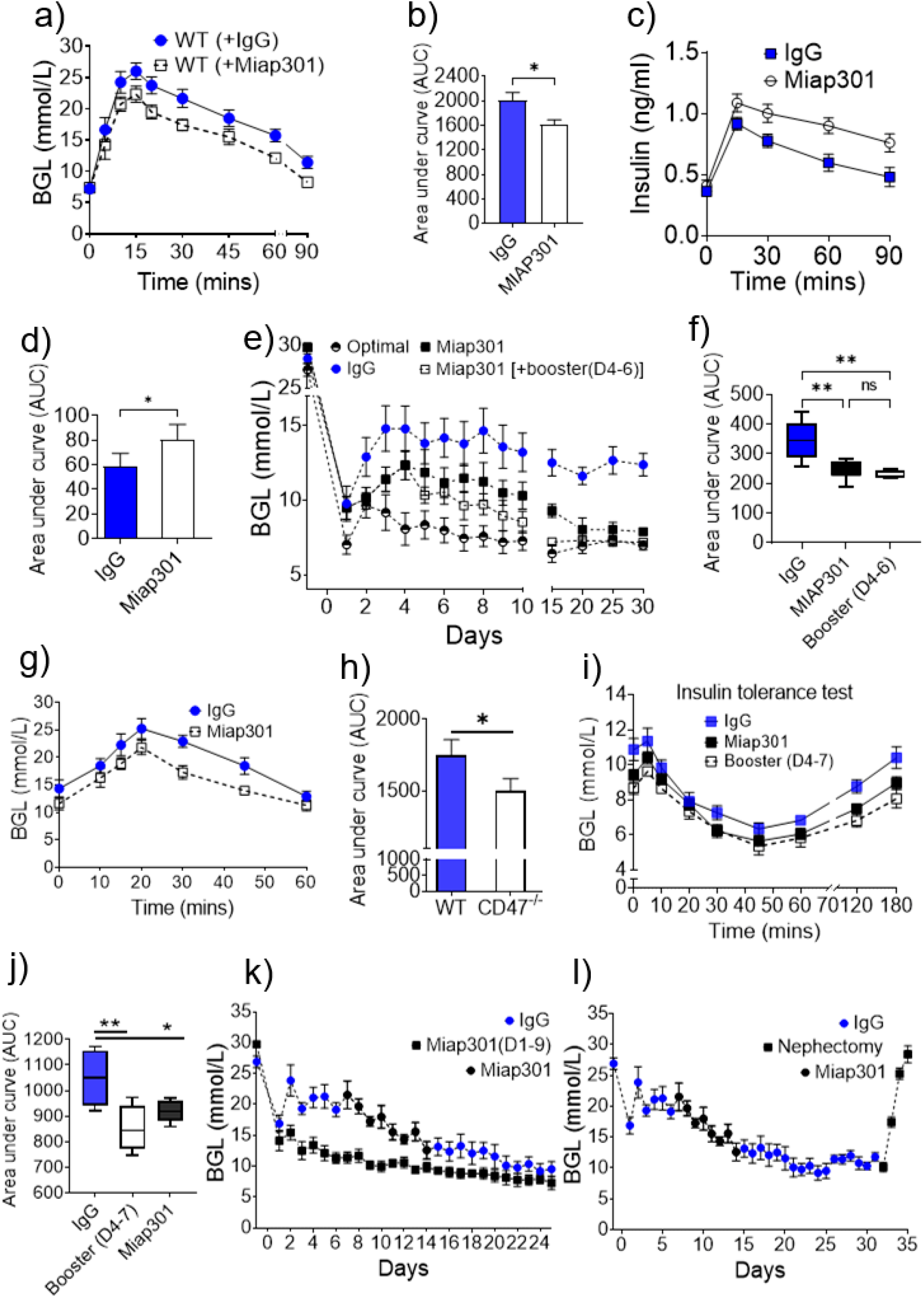
Limited CD47 signaling improves glucose homeostasis and islet transplant outcomes. Twelve-week-old WT mice were treated with IgG (control) or Miap301, 2 h prior to intraperitoneal glucose injection (1 g/kg body weight). (a) Blood glucose levels (BGL) and (b) quantification (area under the curve, AUC) as well as (c) serum insulin levels and (d) quantification (AUC) were measured. e) WT islets were isolated and treated with Miap301 or control IgG for 1 hour before transplantation. Islets were transplanted into diabetic mice (n=6/group) and BGL monitored daily until day 30 post-transplant. In a separate cohort, diabetic mice were transplanted with WT islets treated with Miap301 in addition to Miap301-injected into recipients at days 4-6 post-transplantation (empty squares). f) Results from (e) expressed as AUC. (g-h) Glucose tolerance and its quantification (AUC) in IgG and Miap301-treated groups at day 10 post-transplantation. (i-j) Insulin tolerance and quantification (AUC) between IgG and Miap301-treated groups measured at day 30 post-transplantation. k) Diabetic mice were transplanted with WT islets (n = 12 per group) and n=6 mice were treated with Miap301 from day 1-9 (black squares), and the other half were treated with isotype control (IgG, blue circles). Mice originally receiving IgG were then treated with Miap301 (D7-14, black circles) after which treatment was discontinued. Blood glucose levels were monitored until day 31 after which survival nephrectomy was performed (l). Error bars represent the mean ± SEM, *p < 0.05, ** p < 0.01 by 1-way ANOVA followed by Tukey’s multiple comparisons test (f, j) or t-test.

To further test whether pharmacological intervention could improve the islet transplant outcomes, we conducted transplantation experiments using 450 WT islets. In our hands, this number is neither sufficient to bring the BGL below 10 mmol (our diabetic threshold) nor does it allow BGL to exceed 20 mmol (overt hyperglycemia). One hour prior to transplantation, we treated the isolated islets with either Miap301 or isotype control Ab (IgG). As a control for islet quality, another group of C57BL/6 mice were transplanted with an optimal islet mass (550 islets). Recipient mice transplanted with IgG (isotype control)-treated islets failed to achieve normoglycemia and remained diabetic (BGL>10 mmol) throughout the course of the experiment (Fig. 6e, f). In contrast, mice transplanted with CD47-Ab-treated islets achieved normoglycemia by day 10 (Fig. 6e, black squares). This suggests that inhibition of donor CD47 receptor signaling prior to transplantation can substantially improve glucose homeostasis.

In the early post-transplantation phase (Day 0-4), Miap301 pre-treatment did not appear to exert any effect on glycemic control, so we examined whether CD47 blockade in recipient mice would shorten the time frame to euglycemia. For this experiment, in addition to Miap301 treatment of islets prior to transplantation, we treated the recipient mice with Miap301 from day 4-7 (Miap301[+booster (D4-7)]). Short-term dosing of the recipient had a noticeable effect with recipient mice achieving normoglycemia at day 6, compared to day 9, and decreased the BGL far more quickly compared to the original Miap301-treated islet group (Fig 6e, empty squares). This suggests that recipient-based CD47 inhibition supports the early function of transplanted islets. However, it did not lead to significant differences in overall glycemic control (as calculated by area under curve, Fig 6f), showing that pre-treatment of β-cells was equally effective. Strikingly, even as the Miap301 treatment was ceased, both groups of mice were able to maintain normoglycemia until day 30 (Fig. 6e). These data suggest that transient inhibition of CD47 signaling in the peri-transplant period is sufficient to improve β-cell function and normalize glucose homeostasis. Consistent with improved glycemic control, glucose tolerance at day 10 post-transplantation was also significantly improved in the Miap301-treated groups compared to controls (Fig. 6g, h). In addition, Miap301-treated groups showed a superior insulin tolerance compared to control group, measured 30 days post-transplantation (Fig. 6i, j).

The effectiveness of additional Miap301 doses (Fig 6e) in achieving glycemic control prompted us to check whether this treatment could be leveraged to lower blood glucose post-transplantation in the absence of ex vivo pre-treatment of islets. To test the time-dependence of CD47 receptor blockade on islet transplantation outcomes, diabetic mice were transplanted with 400 islets and treated with daily injections of Miap301 or isotype control from day 1. After 6 days, the IgG-treated group also received Miap301 treatment. As expected, mice that initially received isotype control did not achieve normal BGL (Fig. 6k, blue circles, D1-6). However, when switched to Miap301 from day 7, these mice showed a significant improvement in glycemic control (Fig. 6k, black circles). By day 20, these mice achieved comparable glycemic control to mice that had been treated with Miap301 from day 1 (Fig. 6k). Significantly, the improvement in glycemic control was maintained even after discontinuation of Miap301 at day 14 (Fig. 6k). To demonstrate that glycemic control was dependent on the islet graft function, a cohort of these mice underwent survival nephrectomy at day 32, and all mice displayed rapid reversal of glycemic control (Fig. 6l). These data indicate that even delayed short-term CD47 receptor intervention can restore normoglycemia in previously non-functioning islet grafts and provide stability to islet graft function.

### CD47 expression increases in pancreatic islets during diabetes

Previously published LC-MS/MS-based quantitative proteomic screen to examine time-resolved phospho-proteome of NOD mice pancreatic cells following their exposure to glucose and promoting insulin secretion demonstrated a trend of increased CD47 expression, from non-diabetic to overtly diabetic mice [20]. Since the MS-based studies were conducted in 3-4 biological replicates per condition, we decided to examine these findings further and monitor CD47 levels during diabetic progression in a cohort of NOD mice. Western analysis showed that CD47 expression significantly increases in pancreatic islets during the course of diabetes (Fig. 7a), suggesting a potential role for CD47 in ²-cell dysfunction in diabetes.

**Figure 7.**
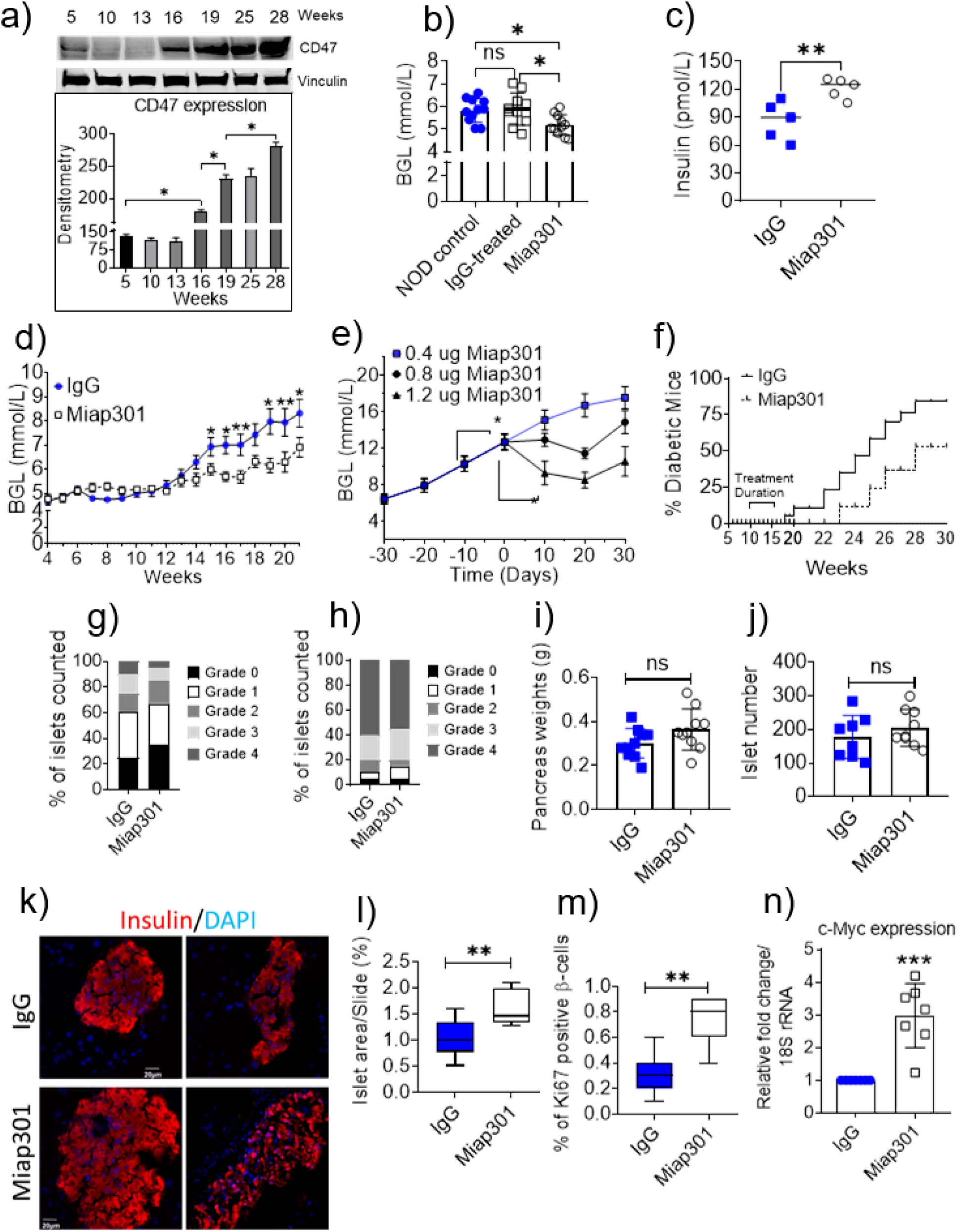
Effects of CD47 blockade on animal model of diabetes. a) Islets from female NOD mice of different ages were pooled and probed for CD47 expression (n=3/age group) and densitometric quantification. b) Blood glucose levels (BGL) and (c) corresponding insulin levels in 6-week old NOD mice following a single dose of Miap301 or IgG (isotype control). (d) 6-week-old female NOD mice were treated with isotype control or Miap301 and BGL monitored (n = 16 per group). e) When blood glucose in Miap301-treated mice reached >10 mmol/L, a 3-fold higher dose of Miap301 was given in an attempt to reverse the hyperglycemia (n=5). f) Percent of NOD mice that became diabetic after either IgG or Miap301 treatment and its time course (n=10). (g-h) Insulitis was scored on islets from NOD mice either treated with Miap301 or isotype control at 15 or 28 weeks of age (n = 5, n = 4 respectively). i) Pancreatic weights of Miap301 vs. IgG-treated WT mice (n=4). j) Total islet number after Miap301 treatment in WT mice (n=4). k) Representative photomicrographs of islets stained for insulin from Miap301 and IgG-treated WT mice (n=4). l) Total islet area of IgG and Miap301-treated WT mice (n=4). m) Quantification of Ki67-positive β-cells in islets from IgG or Miap301-treated WT mice. n) cMyc expression in IgG vs Miap301-treated WT islets. Data are presented as mean ± SEM, *p < 0.05. **p < 0.01. ***p < 001 by unpaired t-test (i-j, l-n) or 1-way ANOVA followed by Tukey’s multiple comparison test.

### CD47 blockade improves glycemic control and delays the onset of type 1 diabetes

Due to the success of Miap301 in controlling BGL following islet transplantation, we considered whether this treatment would also be effective in other settings with reduced insulin secretory capacity. We hypothesized that CD47 receptor antagonism could extend the period of normoglycemia in autoimmune prone non-obese diabetic (NOD) mice. Abnormalities of β-cell function can be detected in NOD mice before they develop diabetes, and this is also true in humans who have dysglycemia before they develop T1D. Following a diagnosis of T1D there is a period of reduced β-cell function that may be sufficient for insulin treatment to be ceased for a period of time (the ‘honeymoon period’). We tested if CD47 blockade would improve glycemic control in NOD mice prior to overt diabetes, with potential application to reducing dysglycemia before and after a diagnosis of diabetes, possibly prolonging the duration of the honeymoon period.

To test the responsiveness of NOD mice to CD47 receptor antagonism, we treated 6-week-old female NOD mice with a single dose of Miap301 before the onset of the dark phase (1900 hours) and measured BGL at 0900 hours the following day. Interestingly, NOD mice treated with Miap301 displayed a significant decrease in glucose levels (Fig. 7b), which was also associated with a trend towards increased insulin levels (Fig. 7c), indicating that β-cells in a T1D-prone environment are responsive to inhibition of CD47 receptor signaling. To determine whether Miap301 treatment would prevent or delay the onset of diabetes in NOD mice, we treated a cohort of NOD mice daily with either Miap301 or IgG from 6 weeks of age. As expected, IgG-treated NOD mice exhibited worsening glucose control over time and eventually developed overt diabetes (Fig. 7d). In contrast, Miap301-treated NOD mice showed a marked resistance to the onset of hyperglycemia for an extended period (3-5 weeks) (Fig. 7d), but eventually these mice treated with standard-dose Miap301 also progressed to hyperglycemia (Fig. 7d). However, when Miap301-treated NOD mice were diagnosed with hyperglycemia (BGL >10 mM), increasing the Miap301 dose 3-fold was able to reverse this trend and further delay the onset of overt hyperglycemia (Fig. 7e). Consistent with the concept that Miap301 treatment improves β-cell function, NOD mice treated with Miap301 showed significantly better glucose tolerance over the time-course of Miap301 treatment at 16 and 20 weeks of age (Suppl. Fig. 3a-d). In comparison, IgG-treated NOD mice showed a worsening glucose tolerance consistent with progression to overt hyperglycemia and diabetes (Suppl. Fig. 3a-d). Next, we determined whether blocking CD47 signaling in β-cells would affect the development of diabetes. We administered Miap301 or IgG to a cohort of 11-week-old euglycemic NOD mice for 5 weeks and monitored blood glucose for the development of diabetes until 30 weeks of age. Remarkably, while 80% of the IgG-treated mice became diabetic by 28 weeks, only 50% of the Miap301-treated mice became diabetic, indicating a significant protective effect of CD47 blockade (Fig 7f). Overall, a short-term blockade of CD47 provided long-term benefit against the onset of diabetes. Investigating the degree of immune destruction in these NOD mice, insulitis scores revealed that overall, there was no significant difference between IgG or Miap301-treated animals at 15 weeks of age (Fig. 7g) and only minor differences into the disease progression at 28 weeks of age (Fig. 7h).

To investigate the general effects of CD47 receptor blockade on islets, we treated 7-week-old WT mice with Miap301 or isotype control (IgG) for 8 weeks. Animals were sacrificed, pancreata isolated, and analyzed. Pancreas weight did not increase significantly in Miap301-treated mice (Fig. 7i), nor was there any difference in total islet numbers between the two groups (Fig. 7j). Islets from Miap301-treated mice appeared to be larger in size (Fig. 7k) and total islet area due to increased individual islet sizes was significantly greater in the Miap301-treated group (Fig. 7l). This increased islet size was most likely due to increased β-cell proliferation consistent with the increased number of Ki67-positive cells seen in Miap301-treated group (Fig. 7m). cMyc is a major regulator of cell proliferation in several cell types, including islets [30, 31]. We and others have shown that loss of CD47 permits sustained proliferation of primary murine endothelial cells and CD47 knockdown or blockade acutely increases mRNA levels of c-Myc and other stem cell transcription factors in cells and in vivo [19, 32, 33]. Our results show that Miap301-treated WT islets had upregulated cMyc levels compared to IgG-treated controls (Fig. 7n). To confirm this further, we isolated islets from WT and CD47^-/-^ mice pancreas and compared cMyc transcript levels under normal and hypoxic conditions. CD47^-/-^ islets expressed higher levels of cMyc under normoxia compared to WT. More interestingly, under hypoxic conditions, there was a significant reduction in cMyc transcripts in WT islets, which was abrogated in CD47^-/-^ islets, confirming a beneficial proliferative effect associated with loss of CD47 that could extend to stress conditions (Suppl. Fig 4a).

## DISCUSSION

Loss of β-cell function within the endocrine pancreas leads to an inappropriate response to fluctuating blood glucose and is characteristic of diabetes or failing islet cell transplantation. In the current study we have identified a new signaling pathway that is regulating insulin secretion in mammals. The cell-surface receptor CD47 plays a constitutive and conserved role in β-cell physiology by tonically inhibiting insulin release in response to a glucose load, providing a potential new therapeutic target to optimally control blood glucose levels.

We observed that CD47 expression is high in islets in vivo, compared to the surrounding acinar cells, which hints at a physiologically relevant role in β-cell function. CD47 expression increases during the course of diabetes development in NOD mice. Using both genetic and pharmacological approaches, we revealed that CD47 inhibition increases glucose-stimulated insulin secretion (GSIS) in MIN6 cells, mouse and human islets. Cdc42 has been implicated as a proximal mediator of insulin release through actin remodeling and facilitating insulin granule movement to the plasma membrane [9], and dysfunction of Cdc42 impairs insulin secretion [25]. However, upstream regulators of Cdc42 in β-cells have remained elusive. Deactivation of Cdc42 appears to be the primary mechanism that CD47 employs to limit GSIS, with evidence of a concurrent increase in insulin docking and exocytosis.

We also demonstrated the superior efficacy of murine islets lacking CD47 in normalizing blood glucose levels in a pre-clinical islet transplant model and replicated these results by blocking CD47 signaling in islets prior to transplantation. Treatment was highly effective in the peri-operative and immediate post-transplant period, suggesting that leveraging CD47 signaling to optimize clinical outcomes could be a temporary, rather than permanent, treatment. In further experiments, CD47 blockade was able to delay the onset of overt diabetes in NOD mice, with negligible differences in insulitis scores (compared to isotype control-treated mice), indicating that improved insulin secretion and not modification of inflammation was the primary driver of glucose homeostasis. These data suggest CD47 receptor antagonism may be clinically beneficial in prolonging the honeymoon period in new-onset T1D patients by enhancing insulin secretion. Humanized monoclonal antibodies to CD47 are currently under investigation in NIH-funded Phase I/II clinical trials for malignancy [34], with development of newer generation antibodies that demonstrate fewer off-target effects. Our work provides robust evidence that would support drug repurposing studies targeting CD47 to enhance insulin secretion. This places CD47 in a privileged position to be relevant in type 2 diabetes (as adjunct treatment to sulfonylureas, biguanides or dipeptidyl peptidase 4 inhibitors), early type 1 diabetes (prolonging the honeymoon period), and islet transplantation (increasing peri-operative function of islets).

Our findings overturn the longstanding dogma in the CD47 field that it is a passive cell surface protein only involved in immune-mediated cell responses as a “don’t-eat-me” signal [14]. CD47 has been thought to regulate cell fate where patrolling phagocytes clear cells that fail to express a minimum threshold of the receptor. Our discovery supports already published works from our and other laboratories that CD47 plays an active role in cellular responses [35], and confirms the rapidly expanding interest in establishing CD47 as a therapeutic target for other diseases. The conserved nature of CD47 expression also suggests relevance in emerging therapies for islet transplantation, including stem-cell derived cellular sources. Data presented here also indicate that strategy to protect stem cell-derived cells against immune-mediated destruction by increasing CD47 expression might undermine their functional properties, at least in case of β-cells.

These findings highlight a potential causal role for impaired CD47 receptor signaling in the development of metabolic diseases such as obesity and diabetes. Importantly, we demonstrate that pharmacological manipulation of CD47 receptor signaling may be a useful therapeutic approach to boost β-cell function under conditions where insulin secretion is limited. Our findings have the potential for a direct clinical application in the context of pancreatic islet transplantation, which is emerging as an important therapy to treat T1D, hypoglycemic unawareness, but is limited in its scope due to the poor function of the transplanted islets after engraftment and scarcity of suitable pancreas donors. Clinical islet transplantation does not enjoy the success seen for solid organ transplants, indicating a need for new therapeutic approaches to improve patient outcomes. Currently, most islet transplant recipients require 2-3 islet transplants to achieve a reduction in hypoglycemic events, and robust metabolic control that facilitates insulin independence. Our studies provide a basis for employing CD47 receptor blockade as co-treatment with islet transplantation to significantly reduce the number of donors required per recipient. This could help reduce the number of donors required per recipient and consequently, increase the number of patients who could benefit from islet transplantation.

Targeting CD47 receptor brings the additional advantage of potentially requiring only a short-term treatment, regime as the effect of CD47 receptor antagonism was highly effective in the immediate period post-transplant period, with efficacy that persisted even after antibody administration was ceased and can be stopped after a short period. CD47 receptor blockade may also be applicable to newly emerging therapies of islet transplantation that utilize alternative islet sources, such as stem cell-derived islets and porcine islets.

## CONCLUSION

Overall, we have identified a novel pathway regulated by CD47 that affects β-cell function and islet transplant outcomes. While some of our findings are reported using the well-established β-cell line MIN6, these data are supported through rodent and human islet studies. We acknowledge the limitations of our *in vivo* glucose tolerance and insulin sensitivity experiments given that global knockout mice were used, and future studies will be needed to understand the peripheral effects of CD47 blockade in insulin-sensitive tissues such as liver, fat and muscle. However, the use of CD47-blocking antibodies robustly recapitulated these findings, supporting proof-of-concept. Further studies will elucidate whether additional signaling partners to CD47, such as signal inhibitory regulatory protein (SIRP), integrins or soluble ligands such as thrombospondin, are concomitant requirements for β-cell function.

## MATERIALS AND METHODS

### Animals

All animal studies were performed under protocols approved by approved by the Western Sydney Local Health District Animal Ethics Committee (#4291) and performed in accordance with the Australian code for the care and use of animals for scientific purposes developed by the National Health and Medical Research Council. Male 3-month old C57BL/6 (CD47^+/+^, wild-type, WT) mice and CD47^-/-^ mice (B6.129S7-Cd47^tm1Fpl/J^) crossed to C57BL/6 mice for 15 generations were purchased from Jackson Laboratory (Bar Harbor, ME). Mice were housed at 23–25 °C in the Westmead animal research facility and maintained under 12 h light/dark cycles at 40–60% humidity with *ad libitum* access to regular chow (8% calories from fat; Gordon’s Specialty Stockfeeds, Yanderra, Australia) and water unless otherwise indicated. Female NOD mice were purchased from Bioservices facility at Walter & Eliza Hall Institute of Medical Research (Kew, Victoria, Australia). To promote obesity, mice were fed ad libitum with a high-fat diet containing lard/sucrose (45% calories from fat, based on rodent diet D12451; Research Diets, New Brunswick, NJ, USA).

### RNA extraction and quantitative RT-PCR

RNA extraction and qPCR were performed as described previously [29]. Specific to these studies, total RNA from mouse islets, human islets and MIN6 cells, were isolated using the RNeasy Mini kit (QIAGEN) following the manufacturer’s recommendations. RNA was quantified using a Nanodrop (BioTek Instruments, Winooski, VT, USA). RNA was reverse transcribed using a SensiFast cDNA Synthesis Kit (Bioline, London, United Kingdom). cDNA was amplified in triplicate with SensiFast Probe No-Rox One-Step Kit (Bioline) and gene-specific Taqman primers (Thermo Fisher Scientific) using a CFX384 real-time PCR machine (Bio-Rad). Thermal cycling conditions were 50°C for 2 min and 95°C for 2 min, followed by 40 cycles at 95°C for 15 s and 60°C for 1 min. Data were analyzed using the ΔΔCt method, with expression normalized to the housekeeping gene.

### Glucose-stimulated insulin secretion ex vivo

For mouse islet isolation, pancreas was perfused with cold liberase (Roche) solution via the common bile duct and subsequently pancreata were removed and incubated at 37°C for 16 min. The digestion was stopped by addition of Krebs-Ringer Bicarbonate buffer containing 10% newborn calf serum followed by mechanical disruption of the pancreas and washing of the islets. Islets were further separated from other pancreatic tissue using a Ficoll-Paque PLUS (GE Healthcare) density gradient and hand-picked with a 20 ul pipette under a stereomicroscope. Islets were subsequently incubated in RPMI-1640 medium (containing 10% fetal calf serum, 1x penicillin and streptomycin, HEPES, 5 mM glucose, and 2 mM glutamine) in a 37 °C, 5% CO_2_ incubator for 24 h. GSIS assays were performed as previously described [36, 37]. Briefly, islets were pre-incubated for 1 h in HEPES-buffered KRB containing 0.1% BSA and 2mM glucose. Experiments were performed in triplicate with batches of five islets incubated at 37 °C, 5% CO_2_ for 1 h in 130 μl KRB containing 0.1% BSA, supplemented with 2.8 mM, 16.7 mM glucose, or as indicated in the text. Supernatant was then collected and insulin release was determined by ELISA kit (Crystal Chem, Elk Grove, IL, USA).

### Human islet insulin secretion

Research-consented human islets unsuitable for clinical transplantation were isolated following the standard protocol carried out by the National Islet Transplantation Unit at the Westmead Hospital, University of Sydney [38]. The use of human tissue for research was approved by the human research ethics committee of the Western Sydney Local Health District. Islets were cultured overnight and insulin secretion experiments were performed with about 80 islets in Krebs solution (114mM NaCl, 5 mM KCl, 24mM NaHCO_3_, 1mM MgCl_2_, 2.2mM CaCl_2_, 1mM KH_2_PO_4_, 10mM HEPES, pH 7.4) oxygenated for 15–20 min at RT and equilibrated for 15 min at 37 °C [39]. Testing was performed as for mouse islets.

### Insulin secretion measurement in MIN-6 cells

For insulin secretion experiment, 3 days prior to experiment, MIN-6 cells were seeded at 25 × 10^4^ per well in a six-well plate. The second day, cells were first washed twice with freshly-prepared KRBH solution (140mM NaCl, 5 mM KCl, 2mM NaHCO_3_, 1mM NaH2PO4, 1.2mM MgCl2, 1.5mM CaCl_2_, 2.8mM glucose, 10mM HEPES, and pH 7.4) and pre-incubated in the same solution for 30 min. The cells were then incubated for 30 min in 2.8mM glucose-KRBH, and 16.7mM glucose-KRBH. Supernatants were collected for insulin analysis by ELISA kit (Crystal Chem, Elk Grove, IL, USA) according to the manufacturer’s recommendations.

### Immunofluorescence and morphometry

Immunofluorescence staining was performed as described [16] with some cell-specific modifications. In short, MIN-6 cells or islets (human and mouse) were seeded onto 13-mm glass coverslips and incubated overnight. At 90% confluence, media was removed, and cells were washed with PBS, fixed with paraformaldehyde (4%) and permeabilized with Triton X at 0.5% in PBS for 10 min. Cells were then washed with PBS and blocked in 5% bovine serum albumin in PBS for 1 h at room temperature (RT), then incubated with the indicated primary antibodies overnight at 4 °C in a humidified chamber. Cells were washed and incubated with Alexa Fluor 488 or 546 secondary antibodies for 1 h at RT. DAPI (Sigma-Aldrich) was used to stain cell nuclei. Cells were washed and mounted in fluorescent mounting media (Dako) and fluorescent images were captured with Olympus FV1000 confocal microscope. For pancreatic tissue or human islets, pptimal cutting temperature compound (OCT compound) was used to embed tissue samples prior to frozen sectioning on a microtome-cryostat. Sections were fixed with 4% PFA for 15 min, then washed and permeabilized for 10 min with 0.5% Triton. Sections were washed and blocked with 2% BSA for 30 min at RT and incubated with respective primary antibodies overnight at 4deg. Sections were washed with PBST and PBS and incubated with Alexa Fluor 488 and 546 secondary antibodies for 1 h at RT. Sections were washed with PBS, then mounted with Vectashield Antifade mounting medium with DAPI, and cover-slipped. Slides were imaged on the Olympus FV1000 Confocal laser scanning microscope. β-cell mass was calculated as described previously [40, 41]. In brief, β-cell ratio assessment was calculated by measuring insulin and acinar areas using Adobe Photoshop in five insulin-stained sections (5 μm) that were 200 μm apart. To calculate β-cell mass, β-cell to acinar ratio was then multiplied by the pancreas wet weight.

### Electron microscopy analysis

For transmission electron microscope (TEM) analysis of mouse islets insulin secretion, mice were anesthetized with tribromoethanol (250 mg/kg) and injected

i.p. with a 2 g/kg bolus of glucose in saline. After 2 min mice were perfused for 30 sec with PBS followed by 5 min with fixative solution (2.5% glutaraldehyde, 0.5% tannic acid, 30mM sucrose in 0.1M cacodylate buffer). The pancreas was then removed and post-fixed for 2 h in the same fixative followed by 1 h in 1% osmium tetroxide buffer. Small, trimmed tissue blocks of about 1 mm3 were washed three times in buffer and stained with 2% uranil acetate in 50% ethyl alcohol for 1 h, dehydrated and embedded in Epon-Araldite resin per standard protocol. Sixty nanometer ultrathin sections were stained with lead citrate and imaged with TEM, (Technai T12 FEI at 1900× magnification). Analysis of cell profile dimension and insulin vesicles content was performed using ImageJ software as previously described [37].

### Glucose and insulin tolerance tests (GTT and ITT)

GTT and ITT were performed as described [19] with some modifications. For the GTT, mice were fasted for 6 h. After fasting, at time 0, a tail cut was made, blood sample taken and fasting blood glucose level (BGL) measured using a Stat Strip Glucometer (NovaBiomedical, Runcorn, UK). A glucose bolus (2 g/kg body weight) was injected intraperitoneally and BGL measured at indicated time points post-injection. For the ITT, mice were fasted for 3 h and fasting BGL measured. Then, an insulin bolus (0.75-1 U/kg body weight) was injected intraperitoneally and BGL measured at indicated timepoints’ post-injection. The area under the curve (AUC) of the glucose measurements was calculated. Serum during GTT was collected and stored for subsequent insulin assays using insulin RIA kits (Linco Research). Fed and fasted serum insulin levels were also measured using insulin RIA kits (Linco Research). Blood samples during GTT were collected and stored for subsequent insulin assays using an ultra-sensitive commercially available ELISA kit (Crystal Chem, Downers Grove, USA).

### Western blotting

SDS-polyacrylamide gel electrophoresis was performed as described previously [42]. Isolated islets and MIN-6 cells were mechanically homogenized in ice cold RIPA lysis buffer (Cell Signaling Technology, Danvers, MA, USA) supplemented with 1x protease inhibitors cocktail (Sigma, St Louis, MO, USA). Samples were centrifuged for 5 minutes at 10,000 g. Protein concentration was determined by BCA (bicinchoninic acid assay) protein assay (Thermo Fisher, Waltham, MA, USA). 20 μg of protein was denatured using Laemmli SDS buffer, heated to 95 °C for 5 min, and subjected to SDS-PAGE. To specifically detect the GTP-loaded forms of Cdc42, the Cdc42 activation kit (Cytoskeleton Inc., Denver, USA) was used according to manufacturer’s instructions. Whole cell lysates (500 μg) were combined with 20 μg of GST-Pak1-PBD-agarose for 24 h at 4°C with constant rotation. After 3 washes with lysis buffer, proteins were eluted from the agarose beads and subjected to electrophoresis on 12% SDS-PAGE. Proteins were transferred to polyvinylidene difluoride (PVDF) membranes. Membranes were immunoblotted with antibody provided in the kit that was specific to Cdc42.

### Islet transplantation

10–12-week-old mice were injected with streptozotocin at doses of 110– 120 mg/kg body weight to induce diabetes. Mice with blood glucose values ≥20 mmol/l were selected as transplant recipients. Transplantation was carried out as previously described [43]. In brief, islets were isolated from pancreas of either CD47^+/+^ (WT) or CD47^-/-^ mice and transplanted into the recipient diabetic mice. For the transplant, the kidney was accessed through a left flank incision. A small nick was made in the kidney capsule at the inferior renal pole and the islets were deposited through the nick toward the superior pole of the kidney. Blood glucose levels were monitored daily. Transplanted mice underwent different regimes as detailed in the text. Nephrectomy was performed at a specific post-operative day as indicated in the text or figures.

### NOD mice treatment plan and study endpoints

Six-week-old female NOD mice received intraperitoneal injections of isotype control (IgG) or anti-CD47 (Miap301) antibody daily (0.4 ug/g body weight). Blood glucose level was monitored for several weeks and when BGL>12mmol was recorded in Miap301-treated, mice were given an increased doses (0.8 ug/g and then 1.2 ug/g) for 30 days to examine the possibility of reverting the trend. In a separate cohort, female NOD mice were treated daily with 0.8 ug/g body weight of Miap301. BGL was monitored weekly prior to 11 weeks of age and twice per week after 11 weeks of age. In these mice IPGTT was carried out at 11-, 16-, and 20-weeks post-treatment. Animals were also monitored daily for signs of deteriorating health, as indicated by weight loss, slow movement, hunched posture, and polyuria. Mice deemed morbid by the above criteria were sacrificed. Pancreas from female NOD mice treated with IgG or Miap301 were formalin fixed, paraffin embedded, and stained with H&E to evaluate islet insulitis. The histology of the pancreatic islets was evaluated by using brightfield microscopy. The insulitis was scored in four levels as previously described [36]. The total number of the islets in the preparations were considered together with the infiltration of lymphocytes. Representations of the different insulitis scores were shown as below: Grade 0 = no insulitis, representing normal islet morphology; Grade 1 = peri-islet insulitis, representing < 25% infiltration with inflammation concentrated around the islet; Grade 2 = intermediate insulitis, representing 25–50% of the islet was penetrated by inflammatory infiltrate; Grade 3 = intra-islet insulitis, representing 50–75% of the islet was penetrated by inflammatory infiltrate; and Grade 4 = complete islet insulitis, representing > 75% of the islet was penetrated by inflammatory infiltrate. The treatments were unknown to the scoring person to ensure blind evaluation.

### Statistical Analysis

Data, unless otherwise indicated, are presented as the mean ± SEM of the results from at least-3 independent cell cultures, 6-8 animals per group, or 3-6 human samples per group. Comparisons were made using the unpaired Student’s t test, and one-way or 2-way ANOVA according to the data type, followed by Tukey’s test for multiple comparisons. A p < 0.05 was considered statistically significant. Statistical analyses were assessed using Prism software (GraphPad Software 6.0 f, Inc, LaJolla, CA, USA). Sample size was predetermined based on the variability observed in prior experiments and on preliminary data. All experiments requiring the use of animals or animals to derive cells were subject to randomization based on litter. Investigators were not blinded to outcome assessment except where otherwise stated.

## Supporting information

Supplemental results

## SUPPLEMENTARY MATERIALS

Figure S1 to S4

## Acknowledgements

The authors thank the imaging facilities at WIMR and Westmead Hospital for their technical support in confocal and electron microscopy. Research-consented human islets isolated at the Westmead Institute for Medical Research were used in the experiments. This work was supported by NHMRC Career Development Fellowship (GNT1158597) to NMR, and Diabetes Australia research grants to KG (DART-G210542) and NMR (DART-19G).

## Author Contributions

All listed authors participated meaningfully in the study. KG and NMR conceived the project and designed the experiments. KG, AK, WJH conducted the experiments. JL and SMJ provided technical support for the study. KG and AK obtained imaging data. KG, AK and NMR analyzed experimental data. STG, JEG and POC provided intellectual input, experimental resources, and protocols for the experiments. KG led the project and wrote the original draft manuscript. KG and NMR edited and revised the manuscript. All authors reviewed and approved the final version.

## Data availability

All data associated with this study are present in the paper or the Supplementary Materials. Data supporting the findings of this study are available from the corresponding author upon reasonable request.

