## Supplemental results for "Novel metabolic role for CD47 in pancreatic β-cell insulin secretion and islet transplant outcomes"

Ghimire *et al.*

1a)

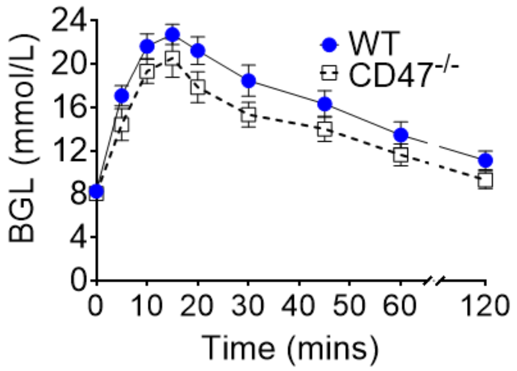


**Supplementary Figure 1: Glucose tolerance test results between 3-month old WT and CD47$^{-/-}$ mice.** a) Intraperitoneal glucose injection (1 g/kg body weight) was performed in WT and CD47$^{-/-}$ mice (n= 8-10 mice). Data are presented as mean ± SEM.

2a)

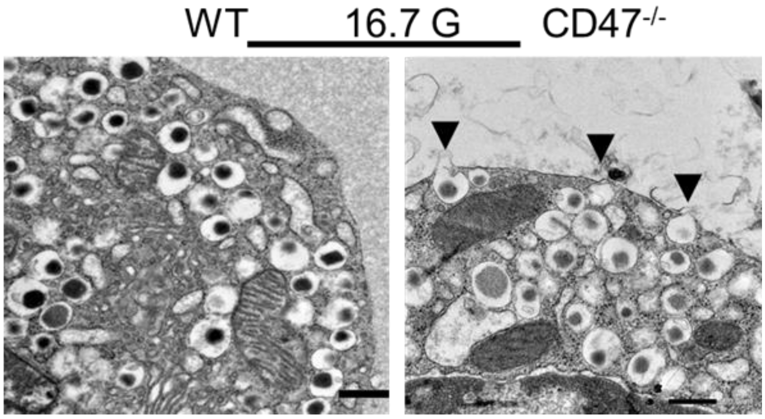


2b)

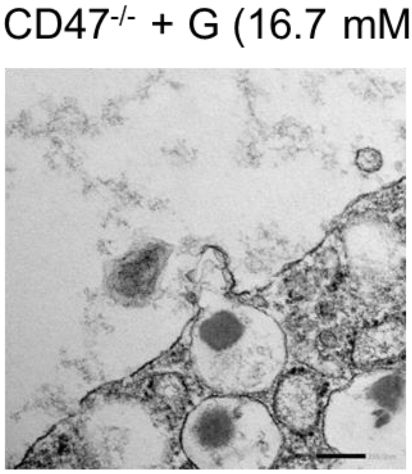


**Supplementary Figure 2: Insulin exocytosis in WT and CD47$^{-/-}$ islets.** a) Representative electron micrograph of exocytosing insulin granules in WT and CD47$^{-/-}$ β-cells following glucose stimulation. Scale bar= 200 nm b) 5x magnification of image in (a) of exocytosing vesicles in CD47$^{-/-}$ islets after glucose stimulation.

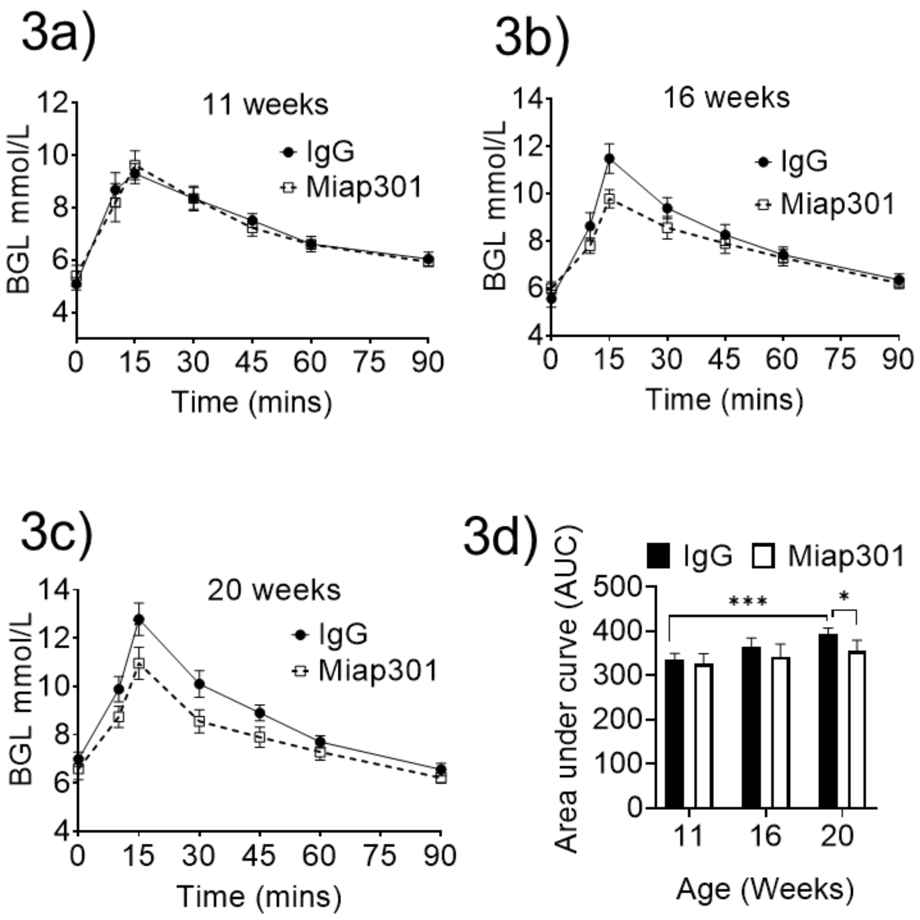


**Supplementary Figure 3: Comparison of glucose tolerance tests between IgG and Miap301-treated NOD mice.** Intraperitoneal glucose injection (1 g/kg body weight) was performed in Miap301- and IgG-treated NOD mice at a) 11 weeks, b) 16 weeks and c) 20 weeks, with (d) quantification (area under curve), n = 6 per group. Data are presented as mean ± SEM, *$p < 0.05$, and *** $p < 0.001$ by 2-way ANOVA followed by Tukey’s multiple comparison test.

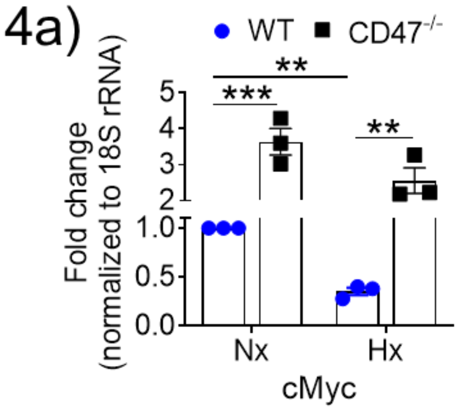


**Supplementary Figure 4: cMyc expression in mice islets under hypoxic stress.** a) cMyc expression in WT and $CD47^{-/-}$ mice islets examined with quantitative real-time PCR. Data are presented as mean ± SEM. *p < 0.05, **p < 0.01 and *** p < 0.001, by 1-way ANOVA followed by Tukey's multiple comparison test
